# APE1 associates with 60S ribosomes and tRNAs and regulates the expression of IGF2BP1

**DOI:** 10.1101/2023.12.08.570814

**Authors:** Wai Ming Li, Belal Tafech, Chow H. Lee

## Abstract

Apurinic/apyrimidinic endonuclease 1 (APE1), a multifunctional protein known for its DNA repair function and redox regulation, is often found overexpressed in cancers. APE1 can be found in the nucleus, cytoplasm and secreted extracellularly. APE1 subcellular distribution in the cytoplasm is frequently reported in various types of cancer but the biological significance remains unknown. In this study, APE1 in the cytoplasm of HepG2 cells was investigated using various techniques including microscopy, differential centrifugation, sucrose gradient fractionation and CL-IP. APE1 was found to associate with 60S ribosomes and tRNAs under native conditions, suggesting it may have a specific function in the translational machinery. Knockdown of APE1 in HepG2 cells led to increased protein expression of IGF2BP1 as well as enhanced HepG2 cell migration, suggesting that APE1 can act as a tumor suppressor in this cell line model of hepatocellular carcinoma. When APE1 was depleted, the translation of a reporter construct containing the 3’UTR of IGF2BP1 was enhanced. This study provides evidence in support of the role of cytoplasmic APE1 in the control of IGF2BP1 protein translation and sheds light on the potential novel function of cytoplasmic APE1.

## Introduction

Apurinic/apyrimidinic endonuclease 1 (APE1) is a multifunctional protein and has been most studied for its role as an endonuclease in various DNA repair pathways (1–4). APE1 is also well-known for its role in redox regulation of transcription factors (5,6). More recently, besides its role in DNA repair, APE1 has gained attention for its ability to process RNA (7–12). APE1 has been shown *in vivo* to be involved in regulating mRNA stability (8), rRNA biogenesis (9), abasic RNA cleavage (7,13), miRNA processing (11,14) and expression (15,16). All these biological functions of APE1 may be attributed to its ability to cleave single-stranded RNA at specific sites as shown by *in-vitro* studies (7,8,17–20).

In clinical settings, APE1 overexpression mostly correlates with poor prognosis for various types of cancer and thus viewed as a protein highly linked with cancer (21,22). Many efforts have been made to inhibit APE1, either by targeting its DNA repair function or its redox function (21). Inhibition of the DNA repair function is thought to sensitize tumor cells to treatments causing DNA damage. Therapies targeting the redox function of APE1 is believed to reduce transcriptional activation of growth-promoting genes. At the present time, the usefulness of all of the available APE1 inhibitors remains to be substantiated in the clinic (21,22). On the contrary, the RNA processing function of APE1 as a therapeutic target has gained little attention.

Within the cell, APE1 has been shown to be mostly distributed in the nucleus, presumably for its major role in DNA repair, redox regulation of transcription factors, rRNA biogenesis, miRNA processing and abasic RNA cleavage in the R-loops. However, numerous studies have also shown cytoplasmic distribution of APE1, both under normal and disease conditions, but the relevance of such observations remains unclear (23–29). Cytoplasmic localization of APE1 may be related to the metabolic state of the cell as it has been shown in highly proliferative cells such as spermatocytes and hepatocytes as well as highly aggressive cancer cells (24,25,27,29). Within the cytoplasm, APE1 can be found in the mitochondria and is involved in mitochondrial DNA damage repair (21,30–32). However, the mitochondria may not be the only organelle where APE1 is localized within the cytoplasm. In recent years, APE1 is also found in the extracellular space (33–35), making serum APE1 (sAPE1) a potential serologic biomarker (22). Extracellular APE1 has been shown in exosomes (35), but the secretion of APE1 as soluble proteins or in microvesicles directly from the cytoplasm has not been ruled out. At present, the biological significance of APE1 secretion remains unknown but appears to be regulated by acetylation of APE1 (34,36,37). The cytoplasm is likely to be the origin of secreted APE1 but unfortunately cytoplasmic APE1 has not been studied in sufficient detail and there is a major gap in our understanding of APE1 in the cytoplasm. We postulate that understanding this pool of APE1 could better define the role of APE1 in cancer in relation to its multiple functions.

Insulin-like growth factor 2 mRNA-binding protein 1 (IGF2BP1) is an RNA-binding oncofetal protein involved in embryogenesis and oncogenesis. As the name implies, it binds to IGF2 mRNA as well as many other oncogenic mRNAs including c-*myc*, CD44 and KRAS (38–44). IGF2BP1 belongs to a small conserved family of RNA-binding proteins which have two RRM domains and four KH-domains for RNA binding. IGF2BP1 binds to target mRNAs to regulate their stability, translation and/or localization (43,44). More recently, the family of IGF2BPs was shown to recognize RNAs in a N6-methyladenosine (m^6^A)-dependent manner and is considered as m^6^A readers (44,45). IGF2BP1 is viewed as a promising target in cancer therapy since many of its targets are involved in cancer growth and development. Notably, IGF2BP1 is particularly known for its role in regulating cell invasion (41,44,46–50) and overexpression of IGF2BP1 is generally correlated with poor prognosis for different types of cancer with a few exceptions (44,51–54). Regulation of IGF2BP1 expression has been demonstrated previously at the transcription and post-transcriptional level mediated by multiple miRNAs including let-7 (44,55).

The present study was carried out with the aim to further our understanding of cytoplasmic APE1. Using microscopy, differential centrifugation, sucrose gradient fractionation and CL-IP, we provide evidence that APE1 interacts with ribosomes and tRNAs under native conditions. Our results also indicate that APE1 can regulate the expression of IGF2BP1 and may have a tumor suppressive role in HepG2 cells.

## Results

### Subcellular distribution of APE1

To determine the subcellular distribution of APE1, we first conducted immunofluorescence studies. As clearly shown, APE1 staining in the nucleus was very strong but upon careful analysis, APE1 was also found throughout the cytoplasm (Fig. 1a) as speckles in HeLa, HepG2, and H441 cells (Fig. 1b and c).

**Figure 1:**
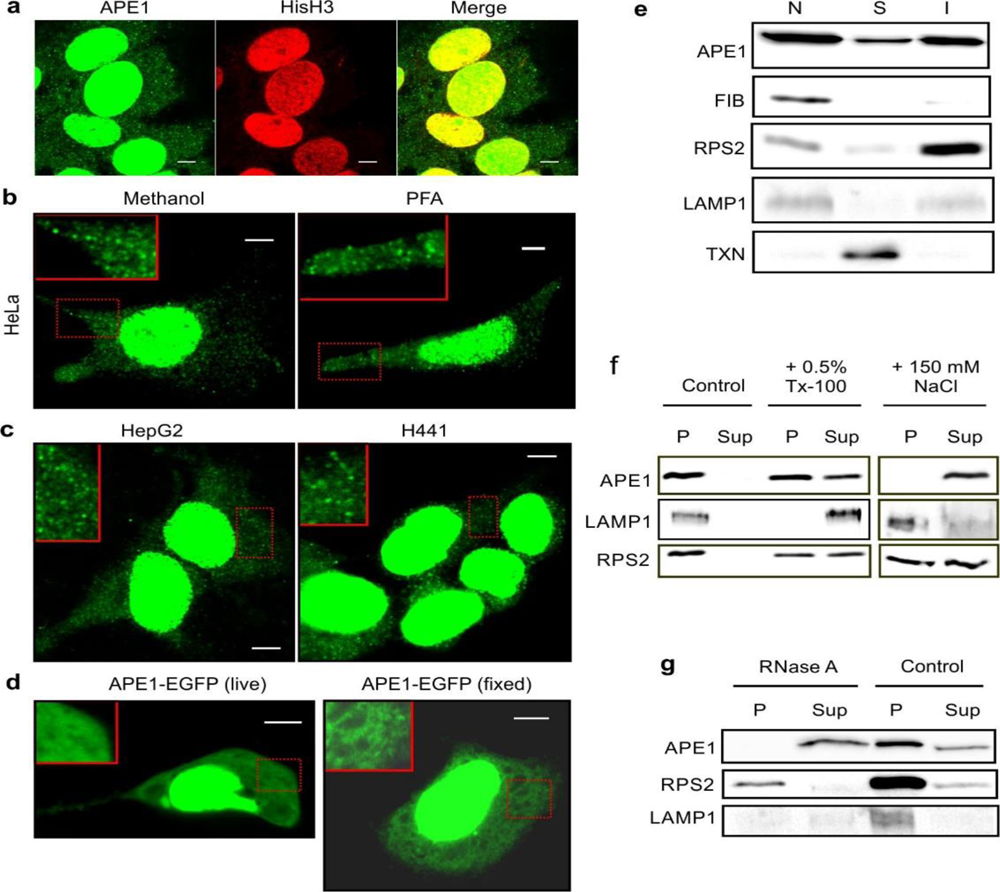
APE1 localizes to both the nucleus and the cytoplasm. APE1 colocalizes with histone H3 (HisH3) in the nucleus in HepG2 cells (a). APE1 is also found in speckles throughout the cytoplasm in HeLa cells (b), HepG2 and H441 cells (c). Different fixation methods were used for immunostaining APE1 in Hela cells (b). Localization of APE1-EGFP fusion protein in live or fixed HepG2 cells (d). APE1 localization in cytoplasmic speckles appears to be abolished when EGFP is attached to the C-terminus of APE1. Scale bar in (a) to (d) = 5 µm. Endogenous APE1 distribution in various fractions from differential centrifugation of HepG2 cells (e). Cytoplasmic APE1 separates into soluble (S) and insoluble (I) fraction after centrifugation at 100,000 x g. Biochemical characterization of fraction I using salt and detergent treatment (f). Cytoplasmic insoluble APE1 (I) obtained from 100,000 x g spin was subjected to treatment with 0.5% Triton-x 100 or 150 mM NaCl for 2 h at 4 °C before re-centrifugation at the same speed to obtain the pellet (P) and supernatant (Sup). Salt treatment (150 mM NaCl) completely removed APE1 from fraction I while APE1 was semi-resistant to detergent extraction. APE1 in fraction I was also sensitive to RNase A (g) as shown by re-distribution of APE1 to the soluble fraction after RNase A treatment and re-centrifugation, suggesting APE1 is associated with RNAs in fraction I. The experiment was performed twice and representative results from one experiment are shown.

The speckled appearance of cytoplasmic APE1 was independent of the fixation method (Fig. 1b). Labelling APE1 at the C-terminus with EGFP appears to abolish the localization of APE1 in cytoplasmic speckles. Diffuse EGFP signal was observed throughout the nucleus and cytoplasm when APE1-EGFP fusion protein was expressed in HepG2 cells when imaged live and after fixation (Fig. 1d). APE1 in cytoplasmic speckles did not appear to colocalize with other known RNA granules such as stress granules, P-bodies and exosomes (Fig. S1), nor with other intracellular organelles and structures such as ER, mitochondria, autophagosomes, endosomes and 40S ribosomes (Fig. S2). However, APE1 appears to be partially colocalized with LAMP1, a lysosome marker (Fig. S2b).

Using differential centrifugation, APE1 was separated into nuclear (N), soluble (S) and insoluble (I) cytoplasmic fractions. Fraction N, which was enriched with the nuclear marker protein fibrillarin (FIB), was obtained in the pellet after centrifugation at 16,000 x g for 30 min. The supernatant was further centrifuged at 100,000 x g to obtain fractions S and I both of which contain APE1. The pellet (fraction I) appears to be enriched in ribosomes containing the marker RPS2 whereas fraction S contains cytosolic proteins like thioredoxin (TXN) (Fig. 1e). APE1 in fraction I was found to be sensitive to extraction using 150 mM NaCl as well as RNase A suggesting that it is associated with structures containing RNA (Fig. 1f and g).

### APE1 associates with 60S ribosomes

To study the subcellular distribution of APE1 in more detail, sucrose gradient fractionation experiments were performed using the supernatant from the spin at 16,000 x g (16K-S) which removed the nuclear fraction. A 7-47% sucrose gradient was used to follow the sedimentation of APE1 under three different conditions to alter the ribosome assembly status. At high Mg^2+^ concentration (1.5 mM) and in the presence of cycloheximide (CHX), ribosomes are prevented from falling off from mRNAs and polysomes were seen in the high density fractions (Fig. 2a, top panel). The majority of APE1 was found in the hydrosol as well as in fractions containing 80S monosomes (Fig. 2a, bottom panel). APE1 was also found to be distributed in polysome fractions while very little was found in 40S fractions. RPL4, which is a marker for the 60S subunit, has a similar distribution with APE1 except in hydrosol fractions. When CHX was omitted in the experiment, less polysomes were present (Fig. 2b, middle panel) as ribosomes are no longer expected to be stabilized on the mRNA. As well, some of the 80S ribosomes became dissociated forming a separate 60S peak (Fig. 2b, middle panel). The majority of APE1 was still found in the hydrosol, 60S and 80S fractions but less in the polysome fractions (Fig. 2b, bottom panel). At low Mg^2+^ concentration (0.25 mM), 80S ribosomes were completely dissociated into 40S and 60S subunits (Fig. 2c, middle panel). Under this condition, APE1 was only found in the hydrosol and 60S fractions (Fig. 2c, bottom panel). The consistent co-sedimentation of APE1 with the 60S subunit under three different conditions demonstrates that APE1 associates with the 60S ribosome whether as a single subunit or in association with the 40S subunit forming mature 80S ribosomes or polysomes.

**Figure 2:**
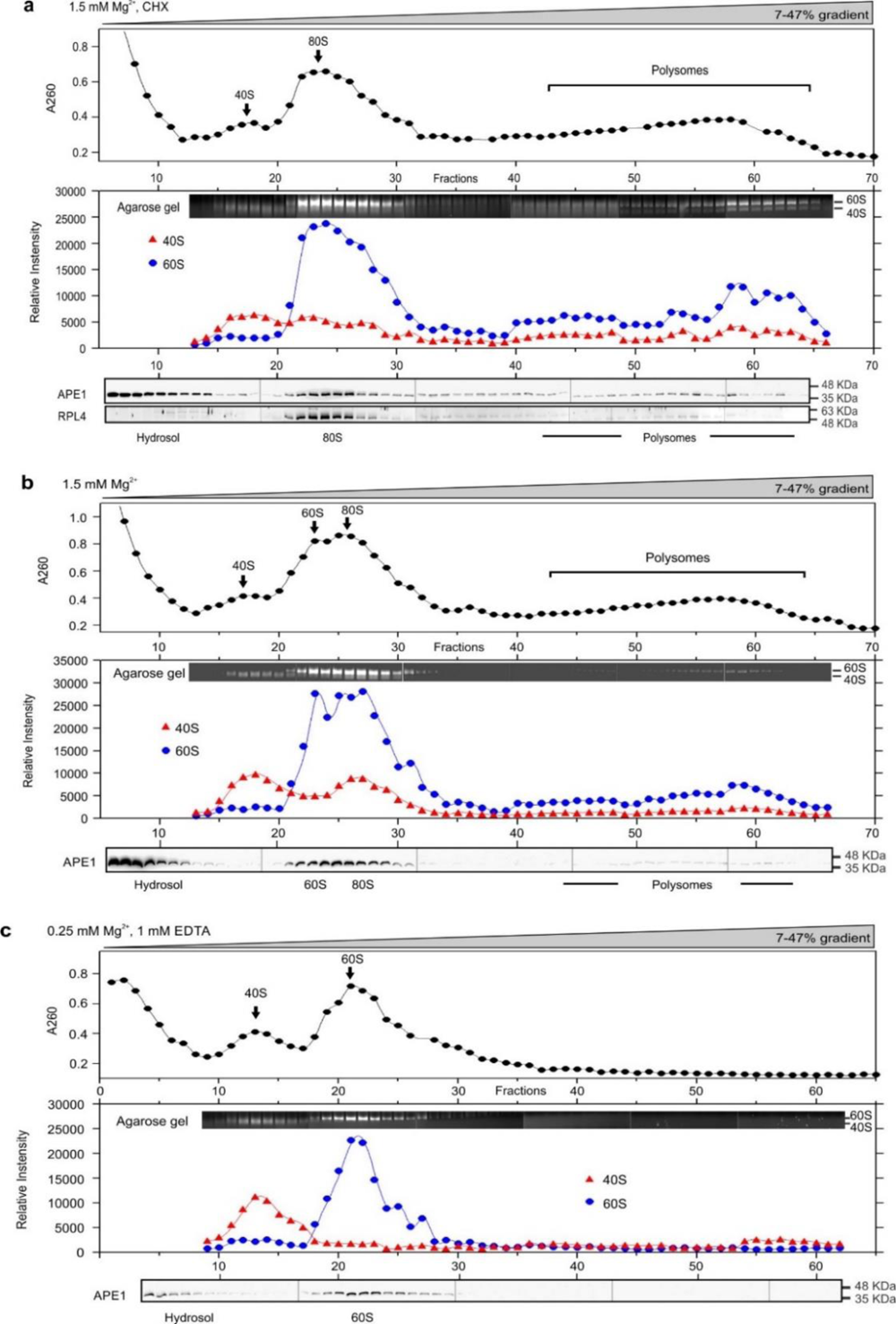
APE1 protein associates with 60S ribosomes as shown by fractionation of cytoplasmic APE1 on a 7-47% continuous sucrose gradient using 3 different conditions to regulate ribosome assembly. (a) At 1.5 mM MgCl_2_ and with 100 µg/ml cycloheximide (CHX) which prevents the disassembly of ribosomes, 40S, 80S and polysomes were observed as detected by A260 (top panel). To detect 40S and 60S subunit separately, fractions were electrophoresed on 1% agarose gel and quantified as relative intensity (middle panel). Western blot detecting APE1 and RPL4 protein in each fraction are shown in the bottom panel. (b) At 1.5 mM MgCl_2_ without CHX, polysomes were diminished and a small 60S subunit peak was observed in addition to the 40S and 80S peak. (c) At low MgCl_2_ (0.25 mM) condition which favors disassembly of ribosomes, only 40S and 60S subunit were observed. Three independent fractionation experiments were performed for each condition and results from one representative experiment are shown. In all conditions, the sedimentation of APE1 in the gradient was associated with the sedimentation of the 60S subunit. APE1 also appeared in the light density fractions (hydrosol fractions) in all three conditions. For each subfigure, total RNA in each fraction was detected by UV at 260nm (top panels) and subsequently separated by electrophoresis on 1% agarose gel to detect 40S and 60S ribosomes (middle panels). APE1 protein in each fraction detected in Western blot is also shown (bottom panels).

### APE1 associates with tRNAs

In addition to the 60S fractions, APE1 was found to be abundant in the hydrosol fractions of the 7-47% gradient (Fig. 2). To further analyze this pool of APE1 which appears to be soluble in the cytosol, a linear 7-27% sucrose gradient was used to better separate the low density fractions containing small RNAs such as tRNAs (∼70 nts). Fractions collected from the top of the sucrose gradient after overnight ultracentrifugation were resolved on 15% Urea PAGE. RNAs of sizes below 100 nts were clearly detected in fractions 23 to 41 with a prominent 70-nt band most likely representing tRNAs (Fig. 3a, top panel). Ribosomes, mostly 40S and 60S subunits (as 80S ribosomes are expected to be completely dissociated at 0.25 mM Mg^2+^), were collected at the bottom of the sucrose gradient and shown as a smear at the top of the gel for fraction 75 (Fig. 3a, top panel). It is interesting to note that the 70-nt band was also detected in this fraction with the ribosomes, confirming that this band is indeed tRNAs which are often bound to 60S ribosomes. Western blot analysis of these fractions shows that APE1 co-sedimented with the 70nt-sRNA with the peak at fraction 27 (black arrow in Fig. 3a, top and middle panel). As expected, APE1 was also detected in the ribosomal fraction at the bottom of the gradient (fraction 75). Alpha-tubulin, on the other hand, known to be a cytosolic protein, sedimented much earlier in low-density fractions with the peak at fraction 17 (black arrow in Fig. 3a, bottom panel). When salt (150 mM NaCl) was included in the experiment (Fig. 3b), APE1 appeared much earlier in the gradient (fraction 17), suggesting that APE1 was released from its binding partner(s) upon salt treatment. Under the same condition, the 70-nt RNA also sedimented in earlier fractions (from peak fraction 27 to 23), suggesting it was also released from its binding partner. It is interesting to note that APE1 in the ribosomal fractions (fraction 75) was also released by NaCl treatment, suggesting APE1 association with 60S ribosomes is also salt-sensitive.

**Figure 3:**
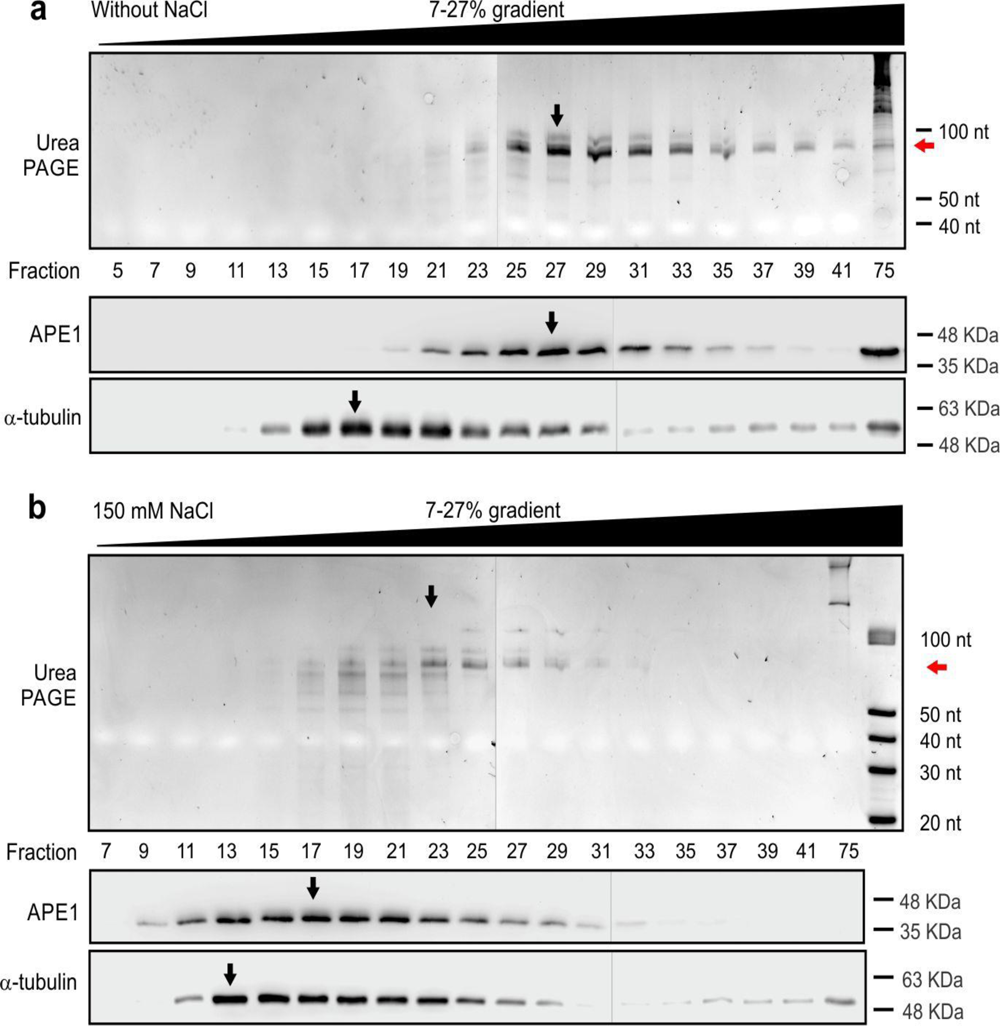
APE1 protein co-sediments with tRNAs. Cytoplasmic APE1 (16K-S) was subjected to fractionation on a 7-27% sucrose gradient to further study APE1 in the low-density fractions without (a) and with 150 mM NaCl (b). Select fractions from the gradient were electrophoresed on 15% polyacrylamide gel with 8M urea to detect small RNAs (top panel). APE1 and α-tubulin in corresponding fractions were detected in Western blot (bottom panel). Red arrows indicate the band which corresponds to the predicted size of tRNAs. Black arrows indicate peak fractions after the relative signal for each band was quantified. The addition of NaCl displaced both APE1 and tRNAs from their binding partners and caused them to sediment earlier in the gradient. The experiment was performed three times for each condition and representative results from one experiment are shown.

To confirm that APE1 is associated with and does not simply co-sediment with tRNAs in the sucrose gradient, *in-vitro* UV cross-linking (CL) experiments were performed. Peak fractions containing both APE1 and tRNAs were collected and then subjected to UV irradiation. After pTex purification to enrich protein-RNA complexes, APE1 was immunoprecipitated and analyzed on Western blot (Fig. 4a). The pTex method was found to be superior to the chloroform/methanol method for enrichment of APE1-RNA complexes which appeared as higher molecular-weight bands as detected on Western blot (Fig. 4a). The small RNAs extracted from immunoprecipitated APE1 showed a strong band corresponding to the size of tRNAs (∼70 nt) when analyzed on 15% polyacrylamide gel with 8 M urea, similar to tRNAs directly extracted from the same fractions used for CL-IP (Fig. 4a, lane 2 and 4, middle panel). When the same fractions from the 7-27% gradient were subjected to IP with APE1 antibody after pTex extraction in the absence of cross-linking, no sRNAs could be detected (Fig. 4a, lane 1, middle panel). When an antibody isotype control was included in CL-IP, no sRNAs were detected (Fig. 4a, lane 3, middle panel). To confirm that the ∼70nt band obtained from CL-IP is tRNAs, Northern blot was performed using probes to detect tRNA^HisGTG^ and tRNA^AspGTC^. Indeed, both of these tRNAs were detected in CL-IP samples when APE1 antibody was used, but not with the isotype control (Fig. 4a, lane 2 and 3, right panel). These results confirm that APE1 in the hydrosol is associated with tRNAs including tRNA^HisGTG^ and tRNA^AspGTC^.

**Figure 4:**
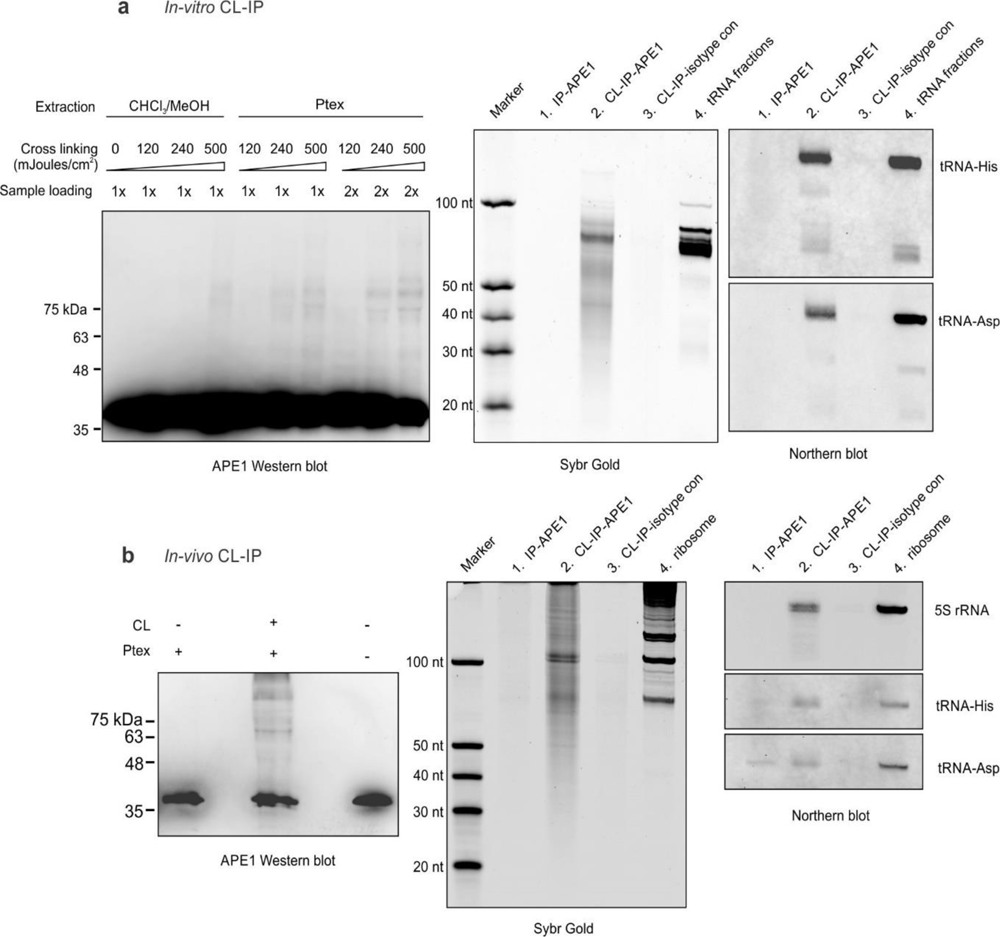
Cross-linking of APE1 to ribosomes and tRNAs. (a) Low density fractions of 7-27% gradient containing tRNAs were UV-irradiated *in vitro* and then purified using pTex method to enrich cross-linked protein-RNA complexes. Detection of APE1-RNA complex by Western blot to compare UV irradiation conditions and protein extraction methods (left panel). Small RNAs extracted after immunoprecipitation of the APE1-RNA complexes were resolved on 8 M urea 15% polyacrylamide gel (middle panel) and was transferred to nylon membrane for Northern blot analysis to detect tRNAs (right panel). (b) HepG2 cells were irradiated to cross-link APE1 to RNAs *in vivo*. APE1-RNA complexes in 16K-S were enriched by pTex method and then analyzed on Western blot (left panel). Small RNAs extracted from APE1-RNA complexes after immunoprecipitation were resolved on 8 M urea 15% polyacrylamide gel (middle panel) and was transferred to nylon membrane for Northern blot analysis to detect and 5S rRNA and tRNAs (right panel). The experiment was performed twice and representative results from one experiment are shown.

### In-vivo cross-linking of APE1 to ribosomes and tRNAs

To demonstrate that APE1 is physically associated with and can be cross-linked to ribosomes and tRNAs *in vivo*, HepG2 cells were irradiated before isolation of 16K-S which contains both soluble and insoluble fractions of APE1. After pTex purification, cross-linked APE1 was analyzed on Western blot (Fig. 4b, left panel). Higher molecular weight complexes of APE1 was observed only in the sample with UV irradiation. Analysis of small RNAs from the CL-IP complexes on 15% urea PAGE again shows a band corresponding to the size of tRNAs (∼70nt) as well as a doublet at ∼120 nt (Fig. 4b, lane 2, middle panel). Northern blot analysis shows that tRNA^HisGTG^ and tRNA^AspGTC^ were detected in the ∼70nt band. As well, the ∼120nt band which appeared as a doublet was determined to be 5S RNA in Northern blot (Fig. 4b, right panel). These results confirm that APE1 binds to tRNAs and ribosomes.

### APE1 depletion enhances the invasiveness of HepG2 cells and IGF2BP1 protein expression

APE1 is overexpressed in many types of cancer including hepatocellular carcinoma (HCC)^21,22^. Higher APE1 expression in HCC is associated with unfavorable prognosis. To test whether APE1 depletion in HepG2 cells can alter its phenotype, we performed knockdown studies using siRNA. We first assessed the growth of HepG2 cells using MTT assay and found that APE1 depletion affected the growth of HepG2 cells minimally (Fig. S3). However, when viewed under light microscope, Si cells looked phenotypically distinct from SN cells (Fig. 5a, top left panels). Long cellular projections were frequently observed in Si cells suggesting these cells may be more invasive. To determine whether that is the case, Si and SN cells were subjected to Matrigel transwell assay (Fig. 5a, bottom left panels) to quantify the number of cells capable of crossing Matrigel and the associated membrane (Fig. 5a, right panel). Indeed, we found HepG2 cells with APE1 depletion were more invasive. Enhanced invasiveness of Si cells could be the result of decreased expression of cell adhesion proteins such as E-cadherin (CDH1). However, we found that CDH1 expression was not significantly different between Si and SN cells (Fig. 5b, right panel). To our surprise, we found that IGF2BP1 expression was up-regulated (Fig. 5b, middle panel) when APE1 protein expression was substantially reduced (Fig. 5b, left panel). Despite the increase in IGF2BP1 protein level, its mRNA level remained unchanged (Fig. 5c, left panel). With increased IGF2BP1, the level of one of its mRNA targets, IGF2, was shown to be significantly reduced (Fig. 5c, right panel).

**Figure 5:**
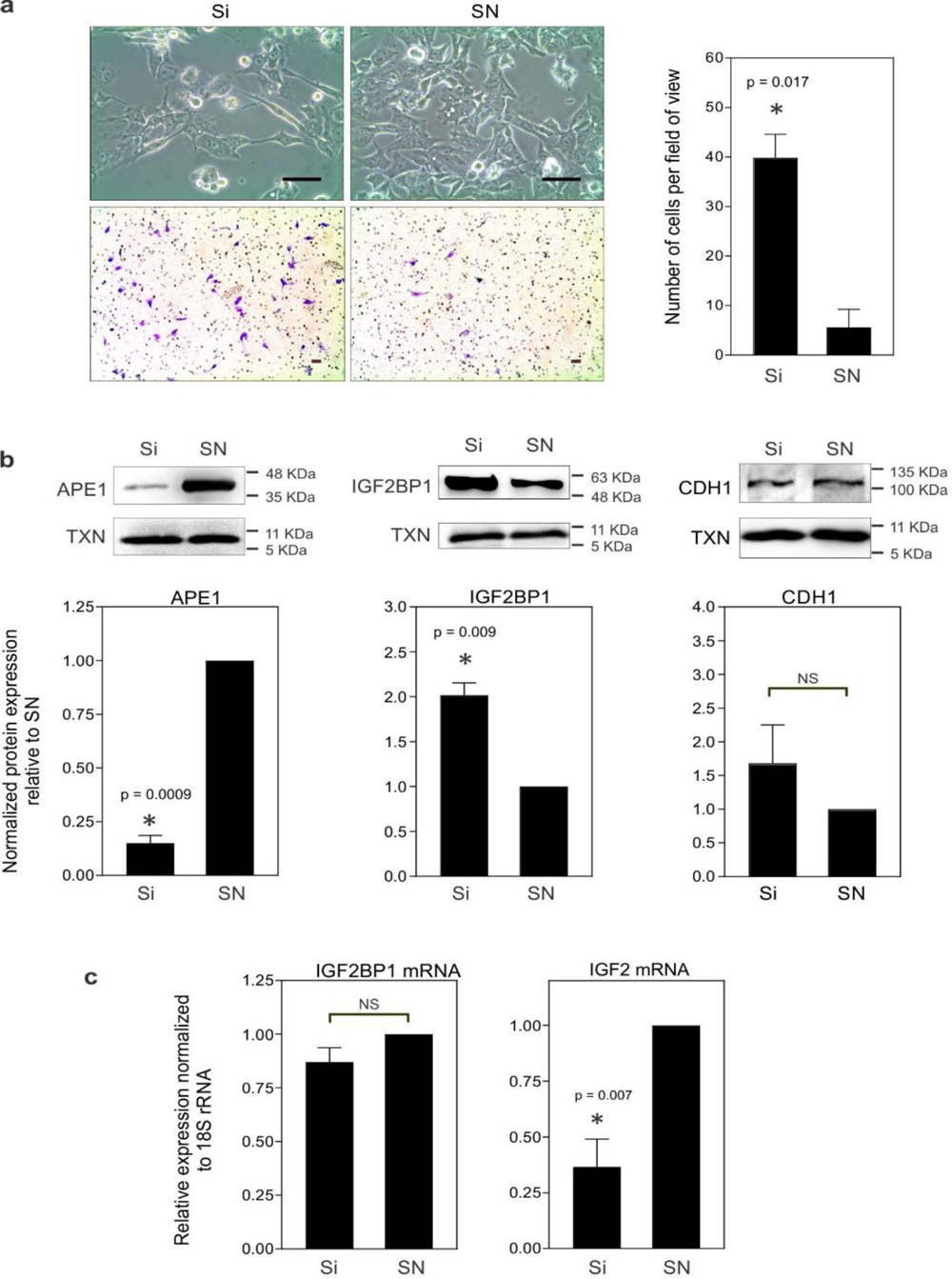
APE1 depletion increases invasiveness of HepG2 cells. (a) Bright-field images of HepG2 cells transfected with APE1 siRNA (Si) and scrambled negative control (SN) (top panels). Matrigel invasion assay was performed to assess the invasiveness of these cells. Two days after transfection, cells were harvested and incubated in Matrigel chambers for 24 h before fixation and staining with crystal violet to capture bright-field images (bottom panels). The number of cells that crossed Matrigel and the membrane was quantified. Results presented in graph represents average from 3 biological replicates ± SEM. Scale bar = 50 µm. (b) APE1 knockdown increases IGF2BP1 protein expression. Representative Western blots showing protein expression of APE1 (left), IGF2BP1 (middle) and CDH1 (right) in Si and SN transfected cells are shown in the top panels. Results in graphs were normalized using thioredoxin expression (THX) as reference and presented as averaged relative expression ± SEM using three biological replicates. (c) Relative expression of IGF2BP1 and IGF2 mRNA in Si and SN transfected cells was normalized using 18S rRNA. Results presented were averaged from 3 biological replicates ± SEM. Statistical analysis was performed using Student’s t-test. * denotes p < 0.05. NS denotes not significant.

### APE1 knockdown increases the translation of IGF2BP1

Increased IGF2BP1 protein level measured in Si cells may be due to reduced clearance of the protein in HepG2 cells. To this end, CHX chase assay was performed. However, no difference in IGF2BP1 levels was found between Si and SN cells (Fig. S4). The translation of IGF2BP1 mRNA has been shown to be regulated by multiple miRNAs in its 3’UTR (44,55). We therefore subcloned the 3’UTR of IGF2BP1 to the 3’ end of renilla luciferase (RL) gene in a reporter plasmid to assess its ability to regulate the translation of RL in cells with and without APE1 depletion. For ease of subcloning, the 3’UTR of IGF2BP1 was divided into four fragments (F1 to F4). We constructed reporter plasmids which contain either the first half (F1 + F2) or the second half (F3 + F4) of IGF2BP1 3’UTR as shown in the schematic of Fig. 6a. Inclusion of the second half of the 3’UTR of IGF2BP1 significantly increased the expression of RL in APE1 knocked-down cells (Fig. 6a). To determine whether increased translation of IGF2BP1 is due to increased association of IGF2BP1 mRNA with polysomes, polysome profiling experiments were performed using fractions separated on a 7-47% sucrose gradient (Fig. S5). However, our results showed no obvious difference between Si and SN cells. This observation together with the finding that APE1 binds to tRNAs and 60S ribosomes suggest that APE1 may play a role in controlling the movement of ribosomes on the mRNA transcript during translation elongation. To further assess whether APE1 is involved in controlling the read through of pause sites on mRNA transcripts, a signature ribosome pausing sequence, *XBP1u*, was used. When dual luciferase reporters containing the *XBP1u* sequence or its control (SBP) were transfected into Si and SN cells, no significant difference was found in the translation of the reporter (Fig. 6b), suggesting APE1 could not alleviate ribosome collision caused by *XBP1u*.

**Figure 6:**
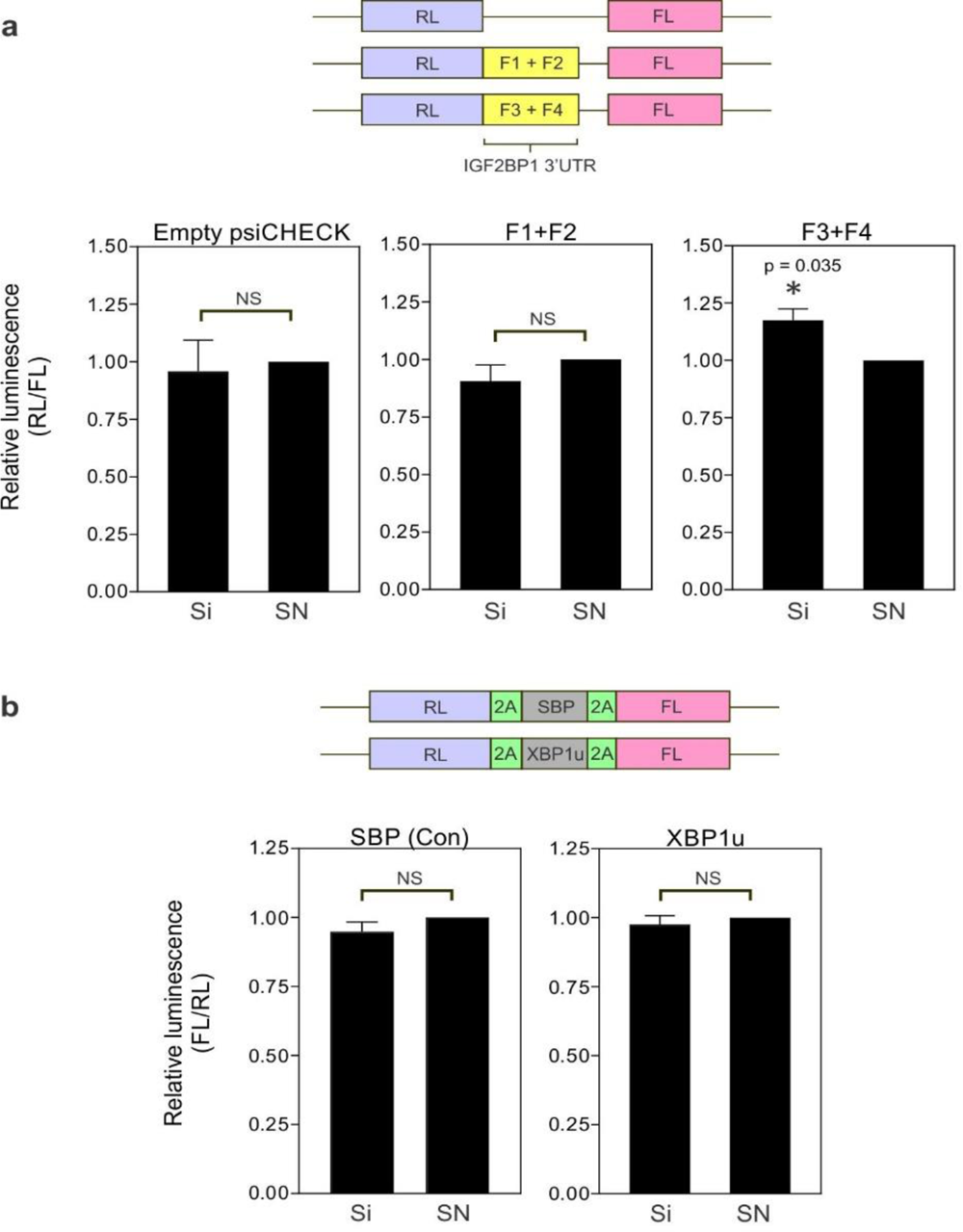
Assessment of protein translation using luciferase reporters in Si and SN transfected cells. (a) Schematic showing reporter plasmids to assess the effect of the 3’UTR of IGF2BP1 on renilla luciferase (RL) translation (top panel). The 3’UTR of IGF2BP1 was divided into 2 halves, with the first half containing F1+F2 and the second half containing F3+F4. Firefly luciferase gene in the plasmid served as a control for transfection efficiency and used for the normalization of RL expression. (b) Schematic representation of the dual luciferase reporter plasmid containing the ribosome stalling sequence *XBP1u* pause site (sandwiched by two 2A self-cleavage sequences) and the control plasmid containing the non-stalling sequence SBP are shown in the top panel. Translational pause due to *XBP1u* site would cause cleavage of the transcript before the FL gene thus reducing the expression of FL relative to RL. Results shown were averaged from three experiments ± SEM. Statistical analysis was performed using Student’s t-test. * denotes p < 0.05. NS denotes not significant.

## Discussion

In this study, we showed that using sucrose gradient fractionation under native conditions, APE1 associates with ribosomes and tRNAs. This newly discovered characteristic of APE1 could explain the many observations of APE1 found in the cytoplasm of various cell types and under disease conditions (23–29). In fact, APE1 was first discovered by us in the ribosomal salt wash of rat liver extracts to cleave c-*myc* mRNA as an endoribonuclease (8,56). Thus, it should be of no surprise that APE1 is found associated with ribosomes. Using a different approach, the 60S ribosomal proteins RPL3 and RPL4 were found to be part of the APE1-protein interactome network although the significance of this finding was unknown (11). Such observation further supports our present findings. In the current study, the association of APE1 to ribosomes and tRNAs was found to be weak because 150 mM NaCl was capable of disrupting the binding (Fig. 1f and 3b). To confirm that APE1 is indeed bound to tRNAs, we performed *in-vitro* CL-IP using sucrose gradient fractions. Two tRNAs, namely tRNA^His^ and tRNA^Asp^, were confirmed to be associated with APE1 (Fig. 4a, right panel). Other tRNAs and their fragments (∼40-50 nt and ∼55-65 nt), which are yet to be identified, may also be bound to APE1 (Fig. 4a, middle panel). By performing *in-vivo* cross-linking experiments, we were able to confirm that APE1 is indeed associated with tRNAs and 5S rRNA which are part of the 60S ribosome (Fig. 4b). We found that immunoprecipitation of APE1 in native conditions (whether in lysates or in sucrose fractions) is impossible without the inclusion of 150 mM NaCl, suggesting the epitope is hidden when APE1 is bound to ribosomes and/or tRNAs. Inclusion of Tx-100 in place of salt improved APE1 IP but no RNAs were found to associate with APE1 unless APE1 was cross-linked before immunoprecipitation. Again, this observation suggests that the RNA-binding site of APE1 is the same site for antibody binding. Another interesting finding from this study is the abolishment of APE1 speckles in the cytoplasm when GFP is fused to the C-terminus of APE1 (Fig. 1d), suggesting the addition of a C-terminal tag may disrupt its normal behavior in the cytoplasm, such as binding to ribosomes and tRNAs. This may possibly explain the different results obtained from IP studies where a FLAG tag was used (11,14). In addition, we believe APE1’s interaction with tRNAs has not been previously reported due to challenges in detecting tRNAs using RNA-seq and also that previous studies did not use specialized protocols for their detection (11).

Our results also demonstrate that APE1 expression is negatively correlated with the protein expression of IGF2BP1 in HepG2 cells and the effect is most likely attributed to post-transcriptional regulation because IGF2BP1 mRNA levels were unchanged (Fig. 5c). One explanation may be that APE1 regulates the level of miRNAs which in turn led to the inhibition of translation of IGF2BP1 mRNA. However, we do not believe this is the case because of the following two reasons. Firstly, the miRNAs which are known to control IGF2BP1 mRNA translation (44) are not regulated by APE1 (11,14). Secondly, our results with increased expression of the reporter containing the second half of IGF2BP1 3’ UTR can only partially account for the increase in IGF2BP1 protein expression (Fig. 5b and 6a). We believe the sequence within the coding region of the IGF2BP1 transcript may also be a contributing factor. Indeed, ribosome collision was shown to be widespread in vertebrates with more than one thousand genes involved (57). In addition to the pause site at the stop codon, IGF2BP1 has two additional sites within the coding region which can cause disome formation (57). Therefore, knowing that APE1 is found on 60S and mature 80S ribosomes, we believe APE1 may have a specific function in ribosomes. What are the possible functions of an RNA cleaving enzyme in the protein translation machinery? Currently, it is believed that the regulation of mRNA metabolism is very much linked to translation, and ribosome is a hub for mRNA decay, quality control and stress signaling (58). One well known translation surveillance pathway is ribosome quality control (RQC) to detect ribosome collision when translation comes to a stall because of the presence of pause sequence in the mRNA transcript. As a consequence, translation stops and the mRNA undergoes no-go decay (NGD). To test whether APE1 is involved in this pathway, we tested whether APE1 depletion would enhance the read through of a transcript containing the well-known pause sequence *XBP1u* (59,60). Using a reporter containing this pause sequence, our results suggest that APE1 knockdown does not alleviate ribosome collision in extreme cases (Fig. 6b). However, this finding does not rule out the possibility that APE1 might be involved in other surveillance mechanisms or even in normal translation. Notably, evidence of widespread co-translational mRNA degradation has been shown in yeast as well as in humans (61,62). Ribothrypsis, the term given to the ribosome-phased endonucleolysis of mRNA, cuts mRNAs at the ribosome exit site thus generating mRNA fragments (62). At the present time, the endoribonuclease responsible for such process remains to be identified. APE1 may have a role in controlling normal translation of certain mRNA transcripts depending on the mRNA sequence.

The finding that APE1 associates with tRNAs also raises many potential roles of APE1 in regulating translation. APE1 has been shown *in vitro* to cleave different RNA targets, including mRNA and miRNAs (7,8,17,18). As well, APE1 can remove the 3’ phosphate group after cleaving RNAs endonucleolytically (20). Small RNAs derived from tRNAs (tsRNAs) are a new class of regulatory non-coding RNAs which includes tRNA halves (tiRNAs) and tRNA-derived fragments (tRFs) (63). The biogenesis and understanding of the biological significance of tsRNAs is currently an active research topic. APE1 may potentially be involved in the biogenesis and recycling of tRNAs after translation, metabolism of tRNAs to generate tsRNAs, or as a tRNA chaperone. Further work is clearly needed to elucidate the significance of APE1’s association with tRNAs.

IGF2BP1 is generally regarded as an oncogene involved in promoting metastasis in many types of cancer. Therefore, its up-regulation, as demonstrated by APE1 depletion in HepG2 cells in this study, is expected to impose a change in the phenotype. Indeed, enhanced transwell migration of HepG2 cells was observed when APE1 was knocked down (Fig. 5a). Such observation raises an intriguing possibility that APE1 can behave as a tumor suppressor, in contrast to the current view that APE1 is an oncogenic protein. In a previous study, secretion of acetylated APE1 can strongly inhibit tumor growth, neovascularization and induce apoptosis after uptake into triple-negative breast cancer xenografts (64). It may be possible that the anti-tumor effect of secreted APE1 is due to translational inhibition after internalization into target cells. Interestingly, angiogenin, another endoribonuclease known to cleave tRNAs, was also demonstrated to be cytotoxic and causes translational repression (65,66). In the cell, perhaps the location of APE1 dictates its function. In the nucleus, APE1 predominantly acts as a DNA repair enzyme as well as a redox factor to activate transcription factors. In the cytoplasm, it could mainly act on RNAs, depending on the availability of the RNAs present. Ribosomes and tRNAs are very abundant in actively dividing cells and naturally become the main targets of APE1. If this is true, it would be very important to design therapeutic strategies to inhibit the right pool of APE1 as a form of cancer treatment.

In conclusion, we have provided evidence in support of the interaction of APE1 with the cell’s translational machinery. The cytoplasm has been traditionally regarded as a mis-localization compartment for APE1 and has been overlooked. Our finding may be the beginning of the discovery of a novel function of APE1 as a translational regulator. Further studies are needed to elucidate the exact function of APE1 with the ribosomes and tRNAs.

## Experimental procedures

### Cells

All cell lines were obtained from American Type Culture Collection and were maintained in Eagle’s Minimal Essential Medium (Lonza) with 10% fetal bovine serum (Life Technologies Inc.) at 37 C in a humidified incubator containing 5% CO2.

### Microscopy

HeLa or HepG2 cells plated on chambered glass slides or cover glass one day prior were fixed in methanol (-20 °C) for 10 min. After blocking for 1 h in block buffer containing 2% BSA and 0.1% Tx100 in PBS pH7.4, the cells were immunostained for APE1 (anti-APE1 C-4, 1:200, Santa Cruz) and another marker using antibodies against calnexin (1:50), COX IV (1:200), RAB5 (1:100), or LC3B (1:200) from a cellular localization antibody sampler kit (Cell Signaling Technology). In other experiments, the cells were immunostained for APE1 and TIA1 (C-20, 1:100, Santa Cruz), DCP1A (1:200, Abcam), LAMP1 (D2D11, 1:100, Cell Signaling Technology), RPS2 (N2C3, 1:200, GeneTex), or DIS3 (1:100, LS-C187155, Life Span BioSciences). Anti-mouse-AF488 (1:200, Molecular Probes) and anti-rabbit-AF647 (1:200, Molecular Probes) were used as secondary antibodies. After each antibody incubation step, the cells were washed 3 times (10 min each) with wash buffer containing 0.1% Tx-100 in PBS. To study the colocalization of APE1 with poly(A)+ RNA, cells were subjected to *in-situ* hybridization after fixation. Hybridization with Cy5-oligo(dT) was carried out at 37 °C overnight in a buffer containing 1 mg/ml yeast tRNA, 0.02% BSA, 10% dextran sulphate and 25% formamide in 2 X SSC. After successive washes with 4x SSC, 2x SSC and then PBS, the cells were subsequently blocked and immunostained for APE1 as described above. Cells fixed in chambered cover glass were imaged immediately. Cells fixed in chambered glass slides were mounted using ProLong® Gold antifade (Molecular Probes) and imaged the next day. All images were taken using an Olympus Fluoview 1000 confocal system with a 60x oil objective (NA: 1.40).

### Differential Centrifugation

Five 100 mm-dishes of HepG2 cells were washed twice with PBS and then harvested in PBS by scrapping. After pelleting the cells by centrifugation at 200 x g, the cells were combined and washed in 600 µl of hypotonic lysis buffer (HLB) containing 10 mM Tris-Cl, pH 7.4, 10 mM KCl, 1.5 mM MgCl2, 1 mM EDTA, 0.1 mM PMSF and 1x Complete protease inhibitor cocktail (Roche, Indianapolis, IN). The cells were allowed to lyse on ice in 900 µl HLB for 45 minutes. To ensure cell separation and more than 90% lysis, the cell suspension was subjected to aspiration 10 times using a 26G needle. After centrifugation at 16,000 x g for 30 min, the pellet was collected and extracted for 1 h at 4 °C using a buffer containing 10 mM Tris-Cl pH 7.4, 420 mM NaCl, 25% glycerol vol/vol, 0.5 mM PMSF, 0.5 mM DTT and 0.2 mM EDTA. The resulting supernatant from another 16,000 x g spin for 5 min was collected as the nuclear fraction (N). The supernatant from the first 16,000 x g spin (16K-S) represents the cytoplasmic fraction and was subjected to ultracentrifugation at 100,000 x g for 30 min using a SW 60 Ti rotor to separate soluble (fraction S) and insoluble APE1 (fraction I) in the cytoplasm. All fractions, N, S and I, were kept at −20 °C until SDS PAGE analysis. Protein content in all fractions was quantified by Bio-Rad Protein Assay Dye Reagent (BIO-RAD, #500-006) and 10-20 µg of proteins were analyzed by Western blot depending on the antigen. All centrifugation steps were carried out at 4 °C.

To characterize cytoplasmic insoluble APE1, fraction I (purified from 9 dishes of HepG2 cells) was subjected to biochemical treatment and was resuspended in 0.5 ml 10 mM Tris-Cl (pH 7.4) containing 1.5 mM MgCl_2_, 0.25 M sucrose and either 150 mM NaCl, or 0.5% Triton-X 100 for 2 h on ice. In a separate experiment, fraction I was subjected to RNase A (1 mg/ml) treatment for 45 min at room temperature. After re-centrifugation at 100,000 x g for 30 min at 4 °C, the pellet and supernatant was collected for SDS-PAGE and western blot analysis.

### SDS PAGE and Western blot

Protein samples from differential centrifugation and sucrose gradient fractionation as well as cell lysates from APE1 knockdown experiments were resolved on 10 or 12% polyacrylamide gel and then transferred onto Amersham^TM^ Protran^TM^ nitrocellulose membrane using a wet transfer system (BIORAD) for 1 h at 4 °C. The blot was blocked for 1 h at room temperature using 5% skim milk in Tris-buffered saline (50mM Tris-Cl, pH 7.4, with 150 mM NaCl) containing 0.1% Tween-20. The blots were incubated overnight at 4 °C with primary antibodies. The following antibodies were used: anti-APE1 (C-4, 1:500, Santa Cruz Biotechnology); anti-fibrillarin (C13C3, 1:1000, Cell Signaling Technology); anti-RPS2 (N2C3, 1:1000, GeneTex); anti-LAMP1 (D2D11, 1:1000, Cell Signaling Technology); anti-TXN (ab26320, 1:2000, Abcam); anti-IGF2BP1 (D9, 1:1000, Santa Cruz Biotechnology); anti-E-Cadherin (24E10, 1:1000, Cell Signaling Technology); anti-RPL4 (NBP1-81329, 1:250, Novus Biologicals); and anti-α-tubulin (ab4074; 1 µg/ml, Abcam). Secondary antibodies used were: anti-rabbit-IgG-HRP (W401B, 1:8000, Promega); anti-mouse-IgG-HRP (W402B, 1:8000, Promega); anti-mouse-IgM-HRP (SC2064, 1:4000, Santa Cruz Biotechnology). Detection of chemiluminescence was carried out using SuperSignal West Pico PLUS (Thermo Scientific) or Pierce^TM^ ECL reagent (Thermo Scientific). Blots were imaged using a ProteinSimple FluorChem Q imaging system.

### Sucrose gradient fractionation

Cytoplasmic APE1 in 16K-S obtained from five 100 mm-dishes described above was collected and stored at −80 °C until the time of fractionation. To separate 40S and 60S subunits, 80S monosomes and polysomes, the 16K supernatant containing 350-550 µg RNA was loaded onto a linear 7-47% sucrose gradient containing 20 mM Tris-Cl (pH 7.4) and 0.25 mM (or 1.5 mM where indicated) Mg^2+^ and centrifuged at 4 °C for 90 min at 36,000 rpm in a SW 41 Ti rotor. To separate small RNAs, the 16K supernatant was loaded onto a linear 7-27% sucrose gradient and centrifuged at 4 °C for 16 h at the same speed using the same rotor. Linear sucrose gradients were generated manually according to the method described previously (67). For 7-47% sucrose gradients, layers of sucrose solution (7%, 17%, 27%, 37% and 47%) were added to the ultracentrifuge tubes and frozen at −80 °C one by one starting with the bottom layer first. For 7-27% gradients, only three layers of sucrose solutions (7%, 17% and 27%) were used. Before ultracentrifugation, frozen sucrose gradients were thawed slowly at 4 °C (for 8 h up to 16 h) to generate linear gradients. Fractions (150 µl) were collected manually from the top of the gradient and transferred to a 96-well UV-STAR® microplate (Grenier Bio-one) for A260 determination using a microplate reader. To visualize 40S and 60S ribosomes, 10 µl of each fraction was electrophoresed on 1% agarose gel stained with ethidium bromide. Relative intensity of the bands corresponding to the 40S and 60S ribosomes were quantified using the FluorChem software. All gels used the same exposure conditions within the same fractionation experiment. For small RNAs, 7.5 µl of the fractions were electrophoresed on 15% polyacrylamide gel containing 8 M urea. Ethidium bromide was used to detect small RNAs after electrophoresis. For protein detection, 140 µl of each fraction was subjected to chloroform-methanol precipitation to remove sucrose before SDS PAGE and Western blot analysis. Western blot exposure condition was normalized between separate blots within the same fractionation experiment using a reference sample. In experiments where cycloheximide (CHX) was used, a freshly made CHX stock solution (10 mg/ml dissolved in H_2_O) was added to dishes of HepG2 cells to make a final concentration of 100 µg/ml. The cells were then incubated at 37 °C for 15 min before harvesting. The same concentration of CHX (100 µg/ml) was included in the lysis buffer HLB. For experiments where 150 mM NaCl was used, solid powder of NaCl was added to the 16K supernatant to make a final concentration of 150 mM NaCl before storage at −80 °C. The subsequent fractionation also used sucrose gradients which included 150 mM NaCl. Fractionation experiments using different conditions were performed 3 times and representative results are shown.

### In-vitro cross-linking

To cross-link sRNAs to APE1 *in vitro*, peak fractions containing tRNAs from four 7-27% sucrose gradients were combined (2.4 ml total) and then transferred to a 96-well plate (60 ul/well) for UV cross-linking using the UV Stratalinker 1800 (Stratagene). The 96-well plate was placed on ice 3.5 cm below the UV source and cross-linked with 500 mjoules/cm^2^. Subsequently, cross-linked RNA-protein complexes were purified using the pTex procedure (68) with modifications. Cross-linked samples were divided into 12 eppendorf tubes with each tube containing 200 µl of the cross-linked sucrose gradient sample mixed with 300 µl of the lysis/binding buffer from the mirVana kit (Invitrogen), 600 µl of acid-phenol: chloroform (mirVana kit) and 200 µl of 1,3-bromochloropropane (BCP). After mixing for 1 min using a vortex mixer (Fisher Brand) at setting 7, the mixture was centrifuged at 20,000 x g for 3 min at 4

°C. Three quarters of the top and bottom layer was removed leaving 100 µl of the interphase behind. Subsequently, 400 µl of H_2_O, 200 µl of ethanol, 400 µl of acid-phenol: chloroform and 200 µl BCP was added to the interphase. After another cycle of mixing and centrifugation (same condition as above) to separate the phases, three quarters of the top and bottom layer was removed again. At last, 9 volumes of ethanol (-20 °C) was added to the interphase to precipitate the RNA-protein complexes at −20 °C overnight. The next day, the precipitated RNA-protein complexes were spun down at 20,000 x g for 30 min at 4 °C, air dried and then reconstituted in IP buffer for immunoprecipitation described below. As a control, the same procedure was performed without UV irradiation but subjected to pTex purification.

### In-vivo cross-linking

HepG2 cells were grown in fourteen 100 mm dishes until 50-70% confluent before *in-vivo* cross-linking. For UV irradiation, the medium was removed, washed once with PBS and replaced with 8 ml of PBS. The cells were irradiated on ice with 400 mjoules/cm^2^ and then subjected to differential centrifugation to obtain the cytoplasmic fraction (16K-S). To purify cytoplasmic RNA-protein complexes, 16K-S collected (2.8 ml total) was subjected to pTex purification in 14 tubes. The resultant precipitated RNA-protein complexes generated from 2 tubes were resuspended in H_2_O for Western blot analysis. The remaining RNA-protein complexes were re-dissolved in 470 µl IP buffer (HLB containing 150 mM NaCl and 0.25% Triton-x 100) and incubated overnight at 4 °C with 30 µl anti-APE1 antibody (AF1044, 0.2 mg/mL) and 100 µl protein-G beads (50% slurry equilibrated with IP buffer). Beads were separated from the supernatant by centrifugation (1000 x g for 1 min at 4 °C) and washed 2x with 750 µl wash buffer (10 mM Tris-Cl, pH 7.4, 150 mM NaCl, 0.25% Triton-x 100 and 2 mM vanadyl ribonucleoside complex). Each wash cycle consists of rotation at 4 °C for 15 min and the beads were recovered by centrifugation to remove the wash buffer. After the third wash cycle with 500 µl wash buffer, the bead slurry was spun down and resuspended in 100 µl wash buffer and digested with proteinase K (Qiagen) at 50 °C for 30 min. Subsequently, the beads were subjected to RNA extraction using the mirVana kit to isolate small RNAs for urea PAGE and Northern blot analysis. Polyclonal goat IgG (Ab 37373, Abcam) was used as CL-IP antibody isotype control. As another control, the same procedure was performed using 14 dishes of HepG2 cells which were not UV irradiated, but were subjected to pTex purification, immunoprecipitation with APE1 antibody, and RNA isolation using mirVana kit.

### Northern Blots

RNA samples from CL-IP experiments were resolved on 8 M 15% urea PAGE and then transferred onto nylon membrane for 30 min at 2.5 mA/cm^2^ using a semi-dry electroblotter. After transfer, RNA was cross-linked onto the membrane using the auto setting with the UV Stratalinker 1800 (Stratagene). The membrane was then incubated for 1 hour with ULTRAhyb™–Oligo Buffer (Invitrogen, AM8663) before hybridization overnight with the probes diluted in the same buffer at a concentration of 5 pmol/ml to detect 5S rRNA and tRNAs. After washing the blot with a quick rinse and for 30 min using 2x SSC with 0.5% SDS, the blot was blocked using Intercept® Blocking Buffer (Li-Cor, 927-70001) for 1 h at room temperature. The blot was then developed using Streptavidin-IRDye 800CW conjugate (Li-Cor, C41209-03) in the dark and imaged after washing 3x with PBS containing 0.1% Tween-20. All incubation steps were performed in a hybridization oven with rotation and the steps before blocking were performed at 42 °C. DNA probes conjugated to biotin at either the 5’ or 3’ end were purchased from IDT Inc. The sequences of the probes used are 5S-probe-47 (CCGACCCTGCTTAGCTTCC-BIO) (69), 5’ tRNA^HisGTG^ (BIO-CAGAGTACTAACCACTATACGATCACGGC) and 3’ tRNA^AspGTC^ (BIO-GTCGGGGAATCGA ACCCCGGTC) (70).

### Matrigel Invasion Assay

HepG2 cells previously transfected 3 times with siRNA and SN control were used in the invasion assay. 2 days after the third round of transfection, 1 x 10^5^ cells were seeded into the inserts of BioCoat Matrigel Invasion Chamber (Becton Dickinson Labware, Mountain View, CA) in EMEM without serum. Before seeding cells, the inserts were pre-soaked (both inside and outside) in EMEM without serum for 2 h in a CO_2_ incubator at 37 °C. After seeding cells, the inserts were placed into 6-well plates containing 2 ml of EMEM with 10% FBS. Serum on the outside of the insert acts as a chemoattractant for the cells to cross the matrigel and the membrane at the bottom of the insert. After incubation in the CO2 incubator at 37 °C for 24 h, the cells that remained in the insert were removed by rubbing the inside of the membrane with a Q-tip. Cells that have crossed Matrigel and on the outside of the membrane were then fixed by soaking the insert in 100% methanol for 10 min at room temperature. The insert was then soaked in 0.5% crystal violet (Sigma) in 70% methanol for 15 min at room temperature to stain the cells. After washing the insert with 500 ml water 3 times, the insert was rubbed again using Q-tips to ensure all cells were removed on the inside. Cells that remained on the outside of the membrane were viewed and counted using a light microscope. At least 4 fields of view were counted and averaged for each biological replicate. Three biological replicates were performed.

### Total RNA extraction and RT-qPCR

Total RNA was extracted from APE1 knocked-down cells (2 wells) using the mirVana kit according to the manufacturer’s protocol. Residual DNA was removed from RNA samples (1 µg of total RNA input) using the DNA Free kit (Ambion Inc.) before cDNA synthesis using the iScript cDNA synthesis kit (Bio-Rad). The following primer sets were used: IGF2BP1 (F: 5’-AACCCTGAGAGGACCATCACT-3’; R: 5’-AGCTGGGAAAAGACCTACAGC-3’); IGF2 (F: 5’- ACACCCTCCAGTTCGTCTGT-3’; R: 5’- GGGGTATCTTGGGGAAGTTGT-3’); 18S (F: 5’-CTGCCCTATCAACTTTCGATGGTAG-3’; R: 5’-CCGTTTCTCAGGCTCCCTCTC-3’); GAPDH (F: 5’-GTCTTCACCACCATGGAGAAG-3’; R: 5’-AGTTGTCATGGATGACCTTGG-3’).

### Luciferase reporter plasmids and transfection of HepG2 cells

The construction of the luciferase reporter plasmids containing IGF2BP1 3’UTR, namely psiCHECK2- F1+F2 and psiCHECK2-F3+F4 used in this study is previously described (71). The ribosome stalling plasmid psiCHECK2-2A-3xFLAG-XBP1u_pause-2A and its control psiCHECK2-2A-3xFLAG-SBP-2A was a gift from Dr. Shintaro Iwasaki (Addgene plasmid # 159583; http://n2t.net/addgene:159583; RRID:Addgene_159583); (Addgene plasmid # 159584; http://n2t.net/addgene:159584; RRID:Addgene_159584).

All transfections were done using Lipofectamine 2000 (Invitrogen) and the reverse transfection method where the transfection mix was added to HepG2 cells in suspension at the time of plating. To knock down APE1 gene, the double-stranded Dicer substrate siRNA directed against APE1 was purchased from IDT Inc. The sense sequence is 5’-r(GUCUGGUACGACUGGAGUACCGG)dCA-3’ and antisense sequence is 5’-r(UGCCGGUACUCCAGUCGUACCAGACCU)-3’. As control, the double-stranded Scrambled Negative (SN) was used. The sense and antisense sequences of SN are: 5’- r(CUUCCUCUCUUUCUCUCCCUUGU)dGA-3’ and 5’-r(UCACAAGGGAGAGAAAGAGAGGAAGGA)-3’. As HepG2 cells are difficult to transfect, all APE1 knockdown experiments were performed using two rounds of siRNA transfections consecutively 48 hours apart, or otherwise stated for the experiment. APE1 knock down was performed in 6-well plates with 20 nM siRNA and 40 x 10^4^ cells/well. For transfection of the luciferase plasmids, 280 ng of plasmid DNA was used with 40 x 10^3^ cells/well in 96-well plates. Three biological replicates were performed for all experiments.

### Dual luciferase assay

Dual-Luciferase Reporter Assay System kit (Promega) was used according to the manufacturer’s protocol. Synergy 2 Multi-Mode Reader (BioTeK) was used for measuring renilla and firefly luciferase signals. Lysates from HepG2 cells transfected with the luciferase plasmids in 96-well plates were transferred to another white 96-well plate before luciferase assay reagents were added. Three biological replicates were run for each experiment.

### Statistical analyses

One-tailed paired Student’s t-tests were performed for statistical analysis. P < 0.05 is considered significantly different.

## Data Availability

All data are contained within the manuscript

## Supporting information

This article contains supporting information

## Acknowledgements

We thank Dr. Shintaro Iwasaki for providing the psiCHECK2-2A-3xFLAG-XBP1u_pause-2A and psiCHECK2-2A-3xFLAG-SBP-2A plasmids.

## Funding and additional information

This work was supported by funding from the NSERC Discovery Grant (grant #: 227158 to C.H.L.), the Canada Foundation for Innovation (grant #: 34711 to C.H.L.) and the BC Knowledge Development Fund (grant #: 103970 to C.H.L.).

