## Supplemental Information for "APE1 associates with 60S ribosomes and tRNAs and regulates the expression of IGF2BP1"

Supplementary data:

Figure S1
Figure S2
Figure S3
Figure S4
Figure S5

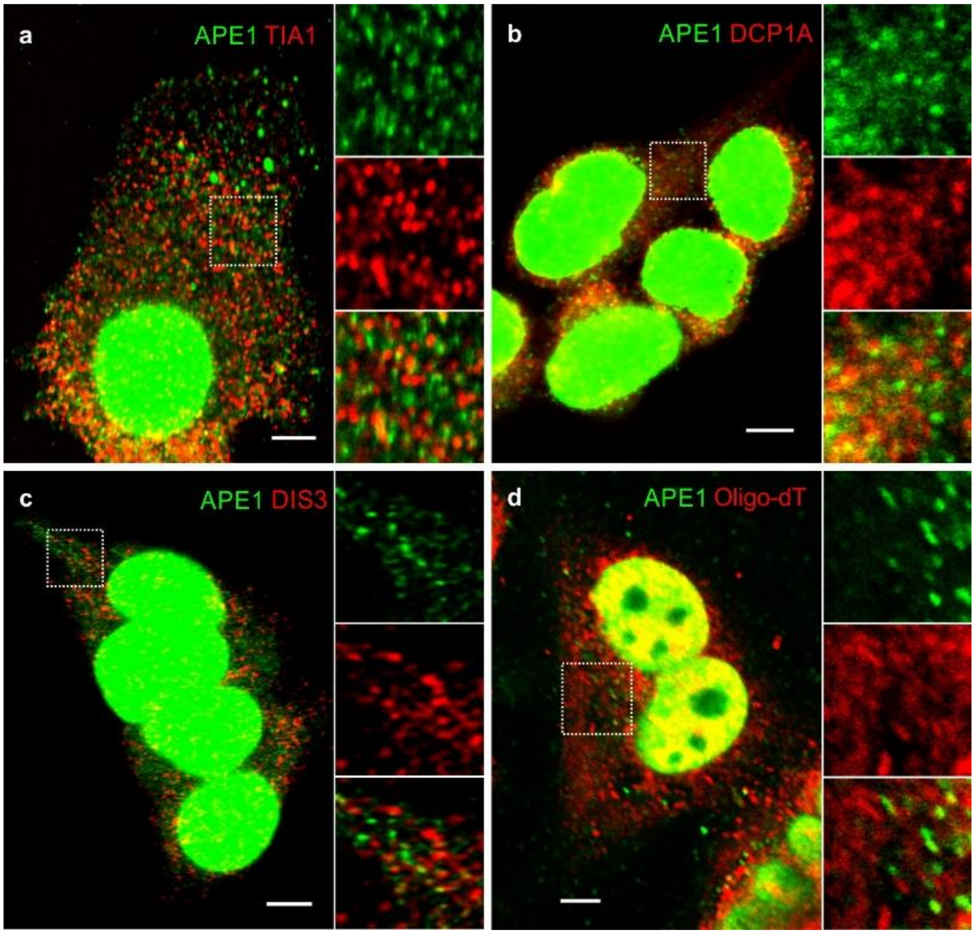


**Figure S1:** Cytoplasmic APE1 speckles do not colocalize with known RNA granules. Stress granules were induced in HepG2 cells with 2 mM $NaAsO_4$ for 3 h before fixation and immunostaining for APE1 and TIA1 (a). H441 cells were fixed and immunostained for APE1 and DCP1A as a marker for P-bodies (b). HepG2 cells were fixed and immuno-stained for APE1 and DIS3 as a marker for exosomes (c). To determine the possible colocalization of APE1 with poly(A)+ RNA, HepG2 cells were subjected to *in situ* hybridization using Cy5-oligo(dT) before immunostaining with anti-APE1 antibody (d). Scale bar = 5 µm.

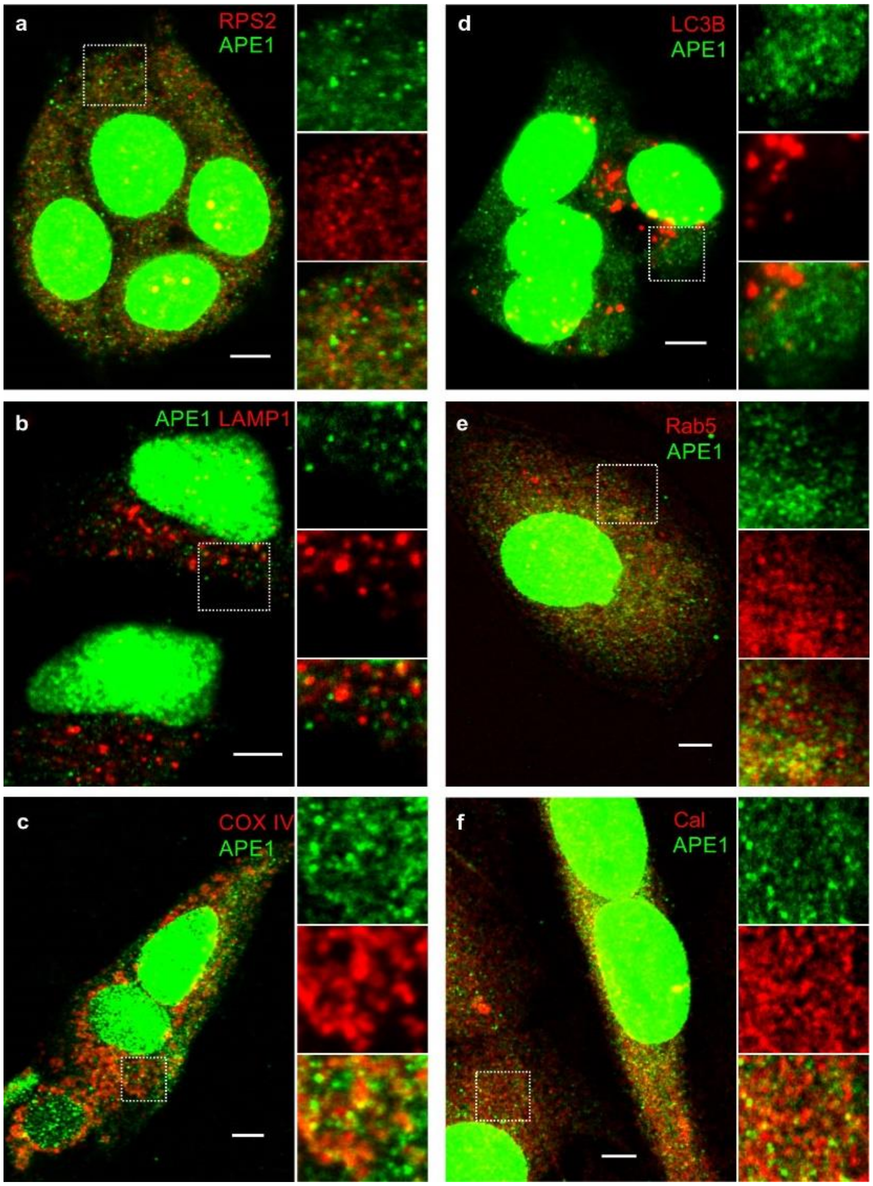


**Figure S2:** Colocalization studies of APE1 with other known intracellular organelles. HeLa (b) or HepG2 (a,c to f) cells were fixed in methanol and then immunostained for APE1 and RPS2 (a, ribosome marker), LAMP1 (b, lysosome marker), COX IV (c, mitochondria marker), LC3B (d, autophagosome marker), Rab5 (e, endosome marker) or Calnexin (f, ER marker) to examine possible colocalization. Scale bar = 5 μm.

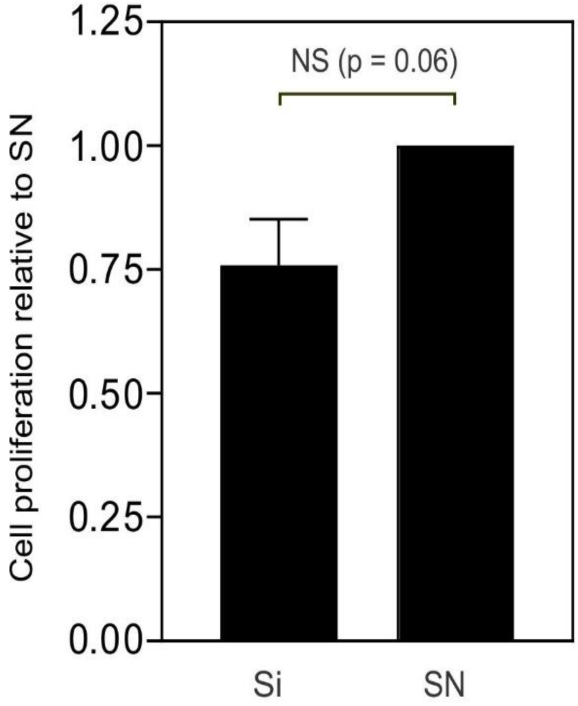


**Figure S3:** Assessment of HepG2 cell growth by MTT assay after APE1 knockdown. HepG2 cells were transfected with siRNA against APE1 (Si) and scrambled negative (SN) twice, 2 days apart. Two days after the last transfection, Si and SN cells were harvested and seeded onto 96-well plates (6000 cells/well). After further culture in a humidified 37 °C incubator for 72 h, MTT assay was performed to assess cell proliferation. Results shown were averaged from three APE1 knockdown experiments ± SEM. Student's t-test was performed for statistical analysis. NS denotes not significant.

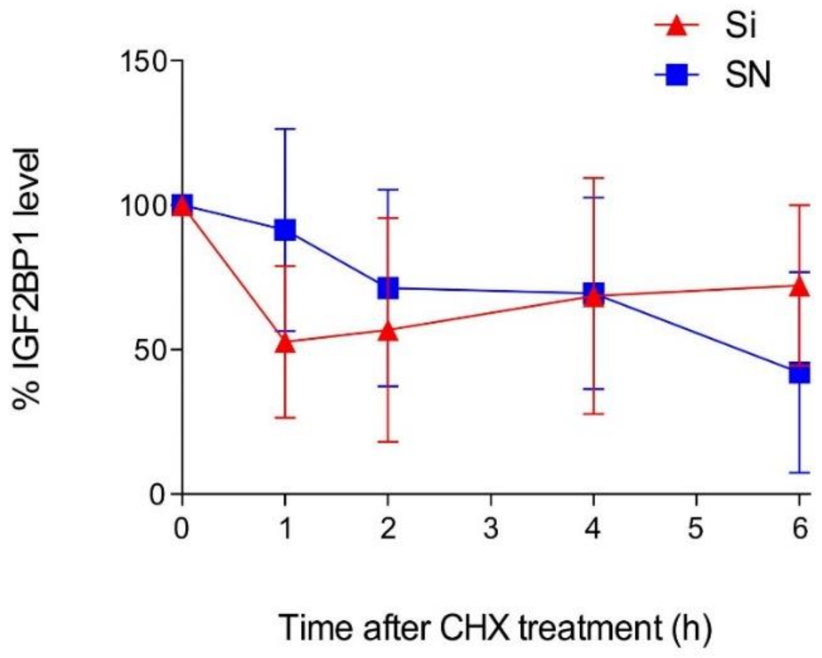


**Figure S4**: Cycloheximide (CHX) chase experiment to assess the clearance of IGF2BP1 protein in APE1 depleted cells. HepG2 cells were transfected with siRNA against APE1 (Si) or scrambled negative (SN) twice, two days apart, before CHX chase experiment. On the day of CHX treatment, medium was removed and replaced with medium containing 300 µg/ml CHX. At 0, 1, 2, 4 and 6 h after CHX treatment, cell lysate was collected for Western blot to quantify IGF2BP1 protein using thioredoxin as the loading control. Results presented were average from three independent APE1 knockdown experiments ± SEM.

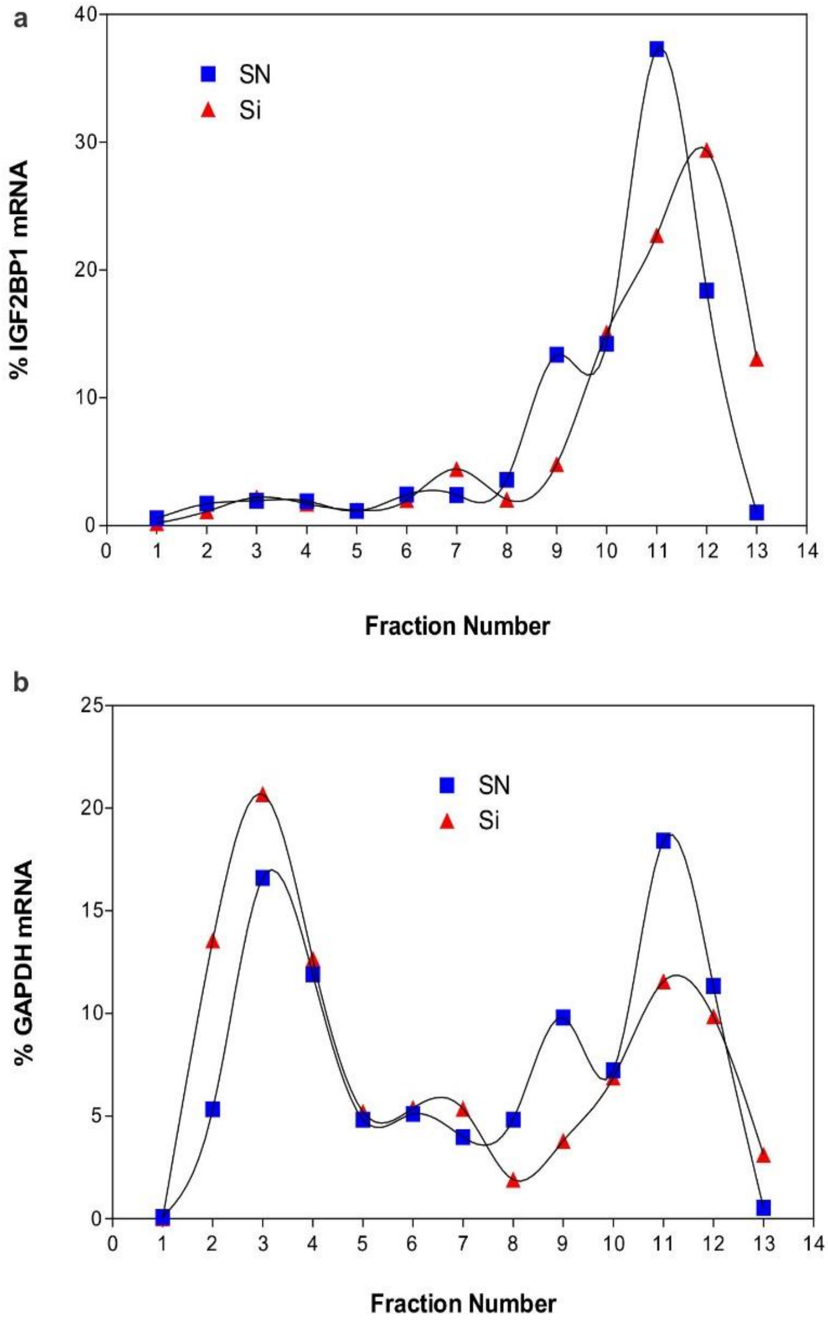


**Figure S5:** Polysome profiling experiment to study the association of mRNAs with ribosomes in a 7-47% sucrose gradient. HepG2 cells transfected with siRNA against APE1 (Si) and scrambled negative (SN) control were harvested to obtain the 16K-S fraction to separate ribosomes on a 7-47% sucrose gradient. The gradient was separated into 13 fractions after ultracentrifugation at 100,000 x g in a SW41 rotor for 90 min and then subjected to RNA extraction using TRIzol reagent. IGF2BP1 mRNA and GAPDH mRNA was quantified by RT-qPCR and expressed as % mRNA in the gradient. The experiment was performed twice and representative results from one experiment were shown.
